# Self-Limiting Factors in Pandemics and Multi-Disease Syndemics

**DOI:** 10.1101/401018

**Authors:** David Manheim

## Abstract

The potential for an infectious disease outbreak that is much worse than those which have been observed in human history, whether engineered or natural, has been the focus of significant concern in biosecurity. Fundamental dynamics of disease spread make such outbreaks much less likely than they first appear. Here we present a slightly modified formulation of the typical SEIR model that illustrates these dynamics more clearly, and shows the unlikely cases where concern may still be warranted. This is then applied to an extreme version of proposed pandemic risk, multi-disease syndemics, to show that (absent much clearer reasons for concern) the suggested dangers are overstated.

The models used in this paper are available here: https://github.com/davidmanheim/Infectious-Disease-Models

## Disease dynamics for extreme events

Dynamics of infectious diseases are complex, and while sophisticated models investigating these factors have been developed, all models simplify various aspects of the spread of a disease. The choice of model (or other decision support tool) depends on the policy or scientific question of interest [15]. Global Catastrophic Biological Risks (GCBRs), discussed by Schoch-Spana et al. [20] are a very extreme class of events, and require somewhat different tools than the ones used in typical epidemiological investigation. Because most infectious disease models are developed and used for typical cases, the usual model assumptions are not appropriate for GCBRs.

Yassif defines a Global catastrophic risk as “something that could permanently alter the trajectory of human civilization in a way that would undermine its long-term potential or, in the most extreme case, threaten its survival.” [23] Given the focus on the overall impact of these events, and their catastrophic impacts, it is particularly critical that models used for assessing that risk appropriately represents asymptotic behavior. Crtically, in an extreme pandemic transmission dynamics will differ from the dynamics appropriate for models which represent more typical cases. Based on conversations with a variety of infectious disease policy and modeling experts [13], this changing transmissibility is the most critical factor identified for assessing the impact of a worst-case pandemic.

The reasons for the expected change in transmissibility are varied, and the dynamics involved in each differ. For instance, social distancing induced by education campaigns [7,11] or self-imposed distancing which occur due to fear would reduce the rate at which contacts occur. Health system responses can lead to isolation of some of the infectious population, or distribute treatments which reduce infectiousness of those infected. Improved hygeinic measures and prophylaxis can reduce the rate at which contacts with infectious individuals lead to infection. In addition, centrally imposed population-level isolation such as border controls and travel screening or restrictions will reduce inter-population spread. Finally, in the most extreme cases, thinning population as the disease spreads limits transmission.

There is an important and large body of work addressing the impact of social distancing on the initial epidemic stages of an outbreak, including information diffusion models [6] and coupled disease-behavior models [22]. This body of work has begun to make use of extant data, however, the asymptotic behavior for human diseases depends on changing dynamics from endogenous response, and these are poorly understood [4]. For example, models of complex heterogenous networks for transmission like those discussed by Pastor-Satorras et al. are critical for understanding initial diffusion [18] but these networks change greatly over the course of a pandemic.

For the purpose of discussing the potential for GBCRs, it is important to choose models that can account for these factors, or adapt models to begin to account for the dynamics important in the extreme cases being discussed. The models mentioned earlier can address these changing population dynamics, but they require detailed data that is impossible to gather without the occurrence of such an event [4,16]. In addition to the data limitation, GCBR cases are likely to come from novel diseases which have unusual and unexpected features [14,20]. Representing the as-yet-unknown features relevant to GCBR cases in detailed models mentioned above would allow modelling the details of spread and transmission across all of the potential cases, but this is infeasible.

Without attempting to represent the dynamics fully, however, simpler models allow exploration of the possibility space. To this end, the paper proposes an adaptation of classical compartmental models that can be used to better explore these limiting dynamics. Such simple compartental models admittedly do not fully capture the dynamics involved, but such simplified models are often able to provide robust results [10].

## Model outline

The proposed model extends the standard SEIR model discussed by Anderson and May [2], by reducing the rate of transmission of the disease as the disease spreads. This is intended to roughly approximate the changing transmission dynamics overall. This allows for clear representation of the disease dynamics without attempting to specify the exact mechanisms involved. Because the various limiting factors are all caused, directly or indirectly, by recognizing the spread of the disease and its seriousness, the changing dynamics are modeled as a function of the cumulative death rate due to the disease. It seems reasonable to presume that at the point where any noticable fraction of a population has died, the population would begin to react to the presence of the disease. For this reason, a linear reduction in the spread of the disease is represented by a reduction in the contact rate. This reduction is expected to continue until a second transition, where the endogenous response is maximal. Beyond this point, the contact rate is reduced linearly until the point of extinction, to represent the reduced transmission rate as population levels drop.

While the compartments are not substantially different than those of a typical SEIR model, to denote the different model assumptions, the model explicitly includes the “Killed” compartment as distinct from “Recovered”, and use the more appropriate term “Latent” for the “Exposed” population. We also explicitly model distancing in the contact rate via the use of *χ* for the contact rate, which depends on fear, *ϕ*, as measured in fatalities. This leads to a Susceptible-Latent-Infectious-Killed-Recovered (SLIKR) model, with a system of ODEs shown below.

**Figure 1.**
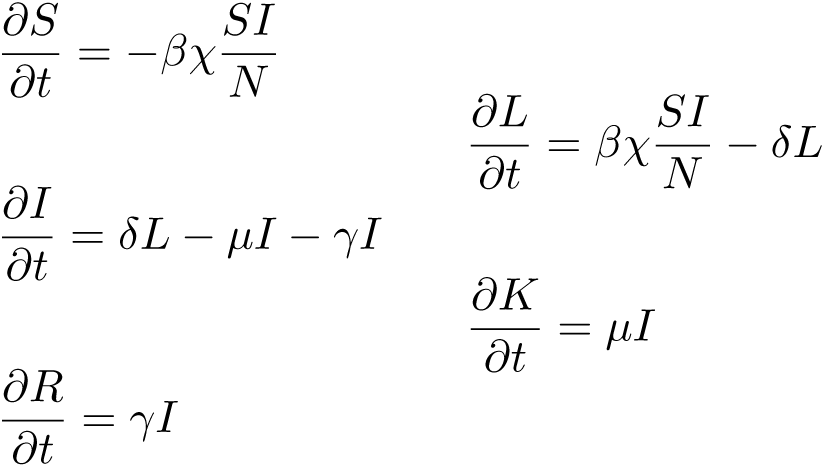
Differential equations for SLIKR dynamics.

N is the total population size. The number of people in each compartment is represented by the compartment initial, and the coefficients are defined below. This formulation assumes a static population.

The contact rate *χ* is non-constant, to represent the changing transmission dynamics. The reduction in contacts is implicitly equivalent to both a reduction in transmission rate, and a reduction in susceptibility, so this variable represents all such changes. The initial level, *χ*_0_, is the normal rate of contacts, transmissibility, and siusceptibility. Once fatalities increase to some minimal level *ϕ*_*min*_, the limiting factors which represent changing public behaviors, health system interventions, begin to have an effect. The limiting factors decrease the contact rate linearly until the death toll reaches *ϕ*_*max*_, at which point the changes have saturated, and infectious contacts are reduced to *χ*_*∞*_. Past this point, we assume contacts decrease linearly to zero at the point of extirpation or extinction, to represent the reduced transmission due to thinned population.

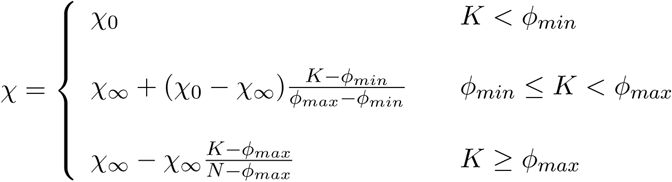

These dynamics are obviously greatly simplified from what will occur in practive, but illustrates the dynamics of social reaction to worst-case pandemics. We also note that the typical alternative of ignoring these dynamics implicitly assumes no distancing occurs, which is even less defensible. To allow exploration of these issues and further elaboration of the model, code for the models used is available on Github, including an interactive model made in RShiny [19] that can be used to explore the dynamics.

One benefit of the models tructure is that the dynamics that differentiate SLIKR models from traditional SEIR models can be zeroed, outputs of the two models can easily be compared. For the illustration of the model, we consider the following dynamics, which are the initial values in the interactive tool. As shown, the final death toll is 12.36%, and the sensitivity to the chosen values can be explored using the interface. If we set *χ*_*∞*_ = *χ*_0_, we can represent typical SEIR dynamics instead, and find that the fatality rate under these assumptions is 70.54%, and we see that representing changing transmission dynamics over time is critical to understanding how a worst-case pandemic unfolds.

| Transformed Variable (Description in tool interface) | Value |
| --- | --- |
| $\beta$ (Probability of transmission per contact) | 5% |
| $\frac{1}{\gamma}$ - (Days Infectious) | 3 |
| $\frac{1}{\delta}$ - (Days Latent) | 2 |
| $\frac{\gamma}{\mu}$ - (P(Death), Case Fatality Rate) | 75% |
| $\chi_0$ - Initial population contact rate | 20 |
| $\chi_\infty$ - Contact rate given maximum response | 5 |
| $\phi_{min}$ - Death rate at which distancing begins | 0.5% |
| $\phi_{max}$ - Death rate at which maximal distancing reached | 2.5% |

## Interactive Tool - Screenshot

### Worst Case Disease Etiologies

The importance and effectiveness of distancing is critical, as shown above. As discussed in an earlier paper [14], this could fail in several distinct ways. First, a disease might spread so robustly that distancing and/or health system responses are ineffective. This would be represented in the model as a reduction in *χ*_*∞*_. If distancing and other interventions are only half as effective as assumed in the example case, *χ*_*∞*_ would be 10 instead of 5, and the death toll in the base case would reach 35.58%, more than double the original 12.36%. Second, a disease might spread acutely, doing so too rapidly for distancing to be effective. If the probability of transmission per contact *β* doubled from the assumed 5% level, the death toll in the example case would reach 41.06%. Lastly, and unfortunately impossible to represent in the current model, the global health system could fail to identify the ongoing spread of a disease. Such a “stealth” disease might have a very long latency, during which it is infectious, or have a post-recovery effect that is only later found to be debilitating.

**Figure 2.**
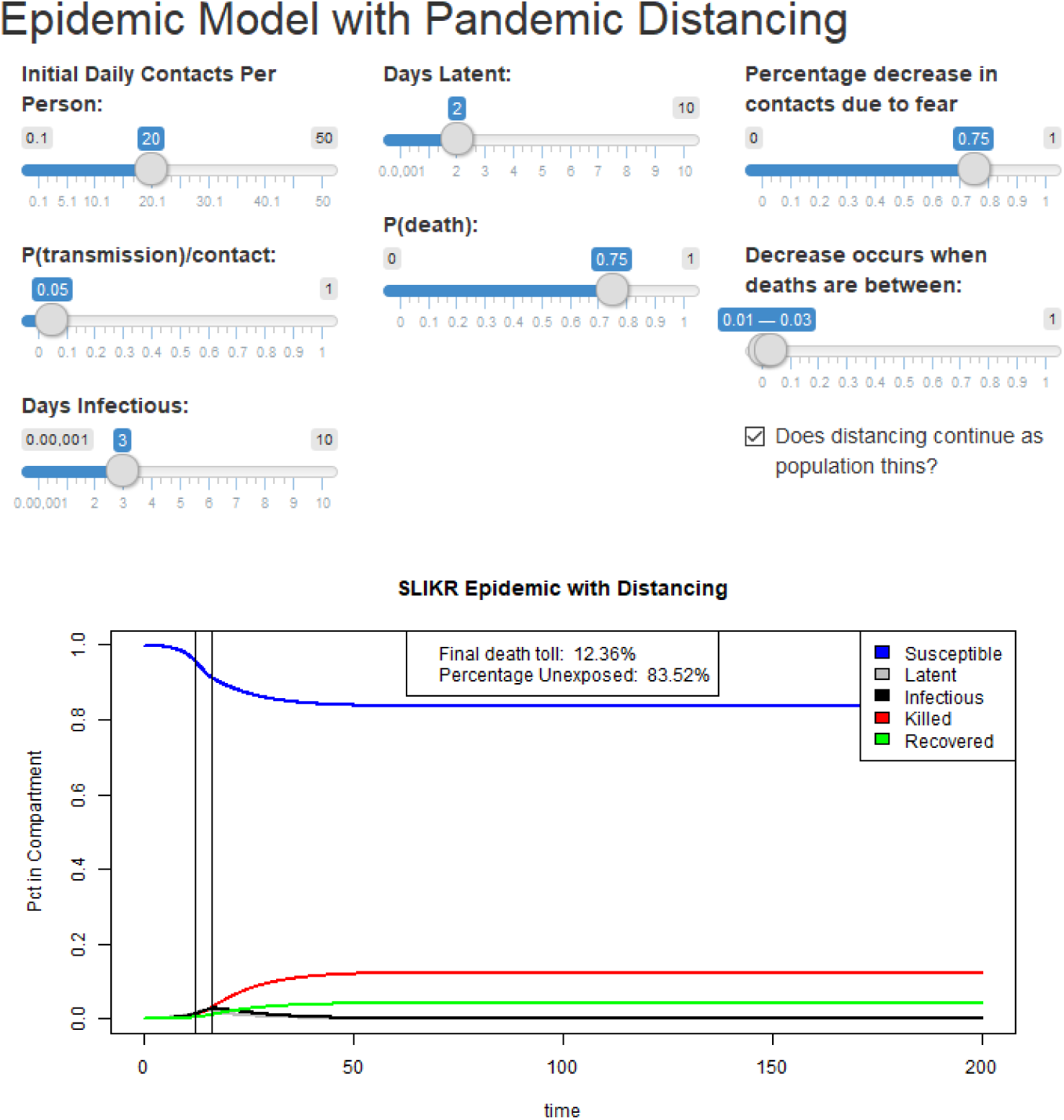
Interactive tool available in the github repository, https://github.com/davidmanheim/Infectious-Disease-Models

There are various other etiologies that have similar effects, such as rapidly spreading diseases among animal reservoirs that can spread to humans, like bubonic plague, which resembles a robust disease, or diseases like AIDS that after a long delay create susceptibility to other infectious diseases, which resmebles a stealth disease. Each of these cases is, of course, similarly not as affected by distancing and is similarly worrisome.

## Model Simplifications

The model simplifications make representing many forms of population response difficult, and instead treat them all as a single continuous phenomenon. This allows easier exploration of simple dynamics, at the expense of the ability to consider different parts of the changing dynamics. In reality, a number of discrete events would be critical, but these are mostly well-represented by the current model structure. For example, while border screening or partial air travel restrictions are unlikely to be effective [3,21], Ferguson et al identify a few discrete policy interventions that are likely to be effective, specifically case-isolation, medical prophylaxis, and reactive school closures [5]. Each of these occurs in a way that is, in general, captured by the current model structure.

On the other hand, some model simplifications are important. As noted, the current model structure cannot represent stealth infections, which is an important phenomenon, as shown by AIDS. The assumption that reaction will be a function only of death rates is also potentially problematic. Because global biosurveillance is effective, health system intereventions could begin much earlier, and warnings to the public could also occur more rapidly. Another key simplification is the linear and permanent nature of the distancing. It is likely that such reactions occur as more of an S-curve, which changes the rate at which people distance themselves and assumes that there is no fatigue or limit to how much people can stay apart. Lastly, the population mixing assumptions inherent in ordinary differential equation models do not allow the model to represent heterogenuous reduction in contacts, which may make distancing much less effective.

## Concurrent Pandemics

An extension of the basic model was developed to consider a question about the potential of existential risks from concurrent pandemics. This has been proposed as a particularly worrying case due to the potential for epidemic-causing bioweapons usage, due either to an accidental release of agents, a concerted attack using several agents, or adversarial use of different agents by different actors. We propose a minimal viable model for addressing the basic question, one which must properly account for disease dynamics of the interactions of multiple diseases. For example, at the very least, such a model must account for the fact that people killed by one disease are no longer spreading other diseases.

There are several distinct ways in which multiple diseases can interact. In some cases, diseases are inhibitory, as with rhinoviruses and other upper respiratory diseases [9], or helminths and microparasites [8]. On the other hand, there is reason to think that syndemics could sometimes have reinforcing effects, as occurs with HIV/AIDs and a variety of other diseases. For this analysis, we follow the assumption in syndemic research outlined in Mustanski et al. that in syndemics, “co-occurring epidemics… additively increase negative health consequences.” [17]

The way in which multiple pandemics would occur and interact will depend on the number of pandemics involved. Unfortunately, compartmental models of syndemics grow rapidly in complexity as the number of simulatneous diseases grow. As shown in Appnedix A, this quickly becomes intractable. The PopMod model developed by Lauer et al [12] models two simultaneous diseases, which can be modeled tractably with our approach, but does not consider infectious diseases, so it is not appropriate for the current application.

The model used in this analysis is capable of representing an arbitrary number of simultaneous diseases that can be fit into a standard SEIR / SLIKR framework, though the computational efficiency of representing multiple models makes the scalability limited. For example, while the interactive model above takes an almost-unnoticeable fraction of a second to update, models with 3, 4, and 5 simultaneous pandemics take 0.5, 2.1, and 9.0 minutes to run respectively. The multi-disease syndemic model has a compartment to represent each distinct combination of Susceptible, Latent, Infectious, and Recovered individuals, and a single compartment for those killed by any of the diseases. (This simplification may be problematic, for example, if there is transmission from corpses like occurs in Ebola. Given the transmission dynamics for the model, however, this possibility is not represented.) The transmission rates and disease progression are assumed to be independent, so that a person with a latent infection of two different diseases will progress to be infectious with one, but retain the latent infection from the other disease.

In order to analyze combinations of different potential diseases, a reasonable but ad-hoc distribution of the disease characteristics was used, and a large number of simulations of the outcomes were run. The distributions used for the disease-specific variables are shown in the table below, while the population characteristics and behavior follows the defaults shown for the single disease model.

| Variable | Distribution |
| --- | --- |
| $\beta$ (Probability of transmission per contact) | Beta(2,20) |
| $\gamma$ - (1/Days Infectious) | Lognormal(1.25,0.5) |
| $\delta$ - (1/Days Latent) | Lognormal(2,1) |
| $\frac{\gamma}{\mu}$ - (P(Death), Case Fatality Rate) | Beta(4,4) |

## Concurrent Pandemic Simulation Results

The simulations were run for 3, 4, and 5 simultaneous disease syndemics, for 1,000, 500, and 100 iterations, respectively^1^. The distribution of the death rate per disease does not differ between the three sets of runs, while the average rate becomes less dispersed as the number of samples increases. It appears that the syndemic death rate, however, is more closely related to the worst single-disease case than to the number of diseases present. This can be understood by considering the interaction between the diseases and the population reaction. Once the public begins to distance themselves due to fear, a syndemic is likely to be slowed and stopped in a fashion similar to what occurs in a single disease. This dynamic is complicated by the interaction between disease spread and death rates.

If one of the diseases spreads more rapidly than the others with a relatively low fatality rate, the population will begin distancing quickly, and the syndemic will be halted before other slower spreading diseases with higher fatality rates have a chance to become established. This can lead to a case where the syndemic death rate is lower than the death rate which would occur due to one or more of the individual diseases. Conversely, if the worst diseases all have similar rates of spread, more than one can gain a foothold before distancing begins. This would allow the syndemic to be worse than any single disease, but in most cases only marginally so. In most cases, however, neither of these dynamics dominate, and we expect a strong relationship between the worst case disease death rate and the syndemic death rate, as seen in the simulations.

To illustrate these dynamics, we can show the population dynamics for two extreme cases found in the 3-disease syndemic dataset, the ones with maximal and minimal difference between the syndemic death rate and the maximum death rate.

In the worst case, distancing started around day 85, and two diseases nearly-simultaneously peaked soon afterwards. These diseases had individual fatality rates of 45.5% and 34.7%, respectively, while the third disease, which was slower to spread, had a fatality rate of 53.3% when it alone was present. In the joint case, as shown in the image, the first two were much more lethal during the syndemic, while the third never spread significantly.

**Figure 3.**
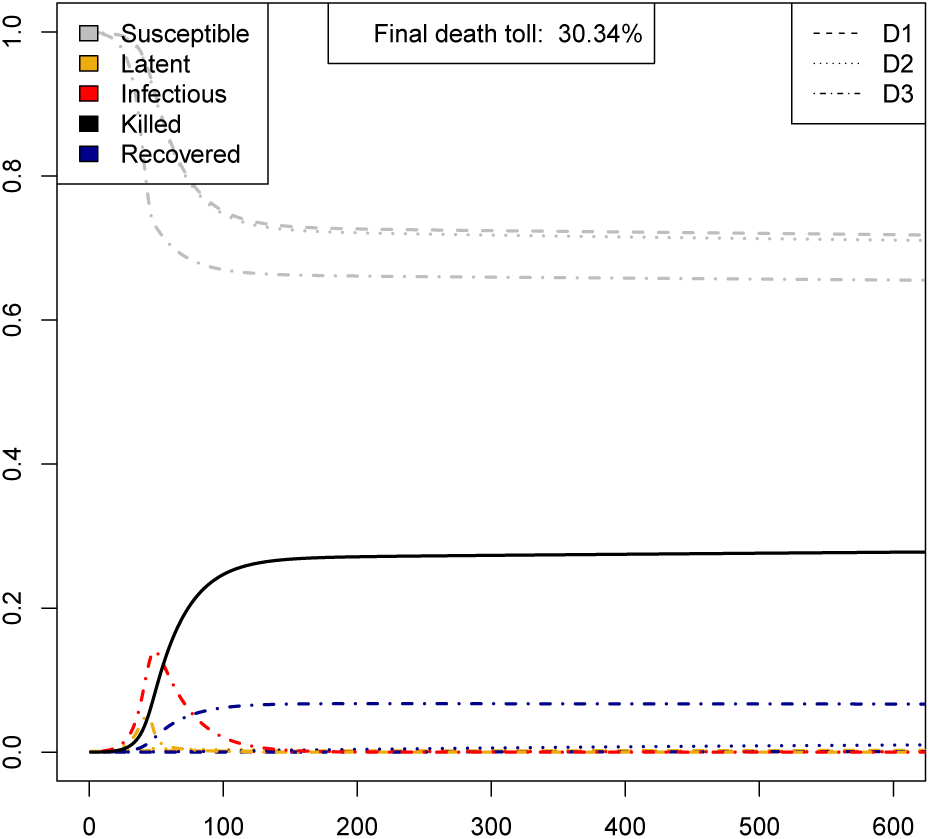
Lowest Syndemic death rate compared to worst individual disease case for 3 simultaneous outbreaks. The figure illustrates the spread of all three diseases when combined in a syndemic. Note that Diseases 1 and 2 are nearly invisible because they are prevented from spreading due to early distancing.

In the least bad case, disease 3 was much more rapid to spread, but had a much lower case fatality rate. For this reason, the overall pandemic death rate, 30.34%, was much closer to the rate for that disease alone, 27.11%, than to the other two less rapidly spreading (but individually worse) diseases.

**Figure 4.**
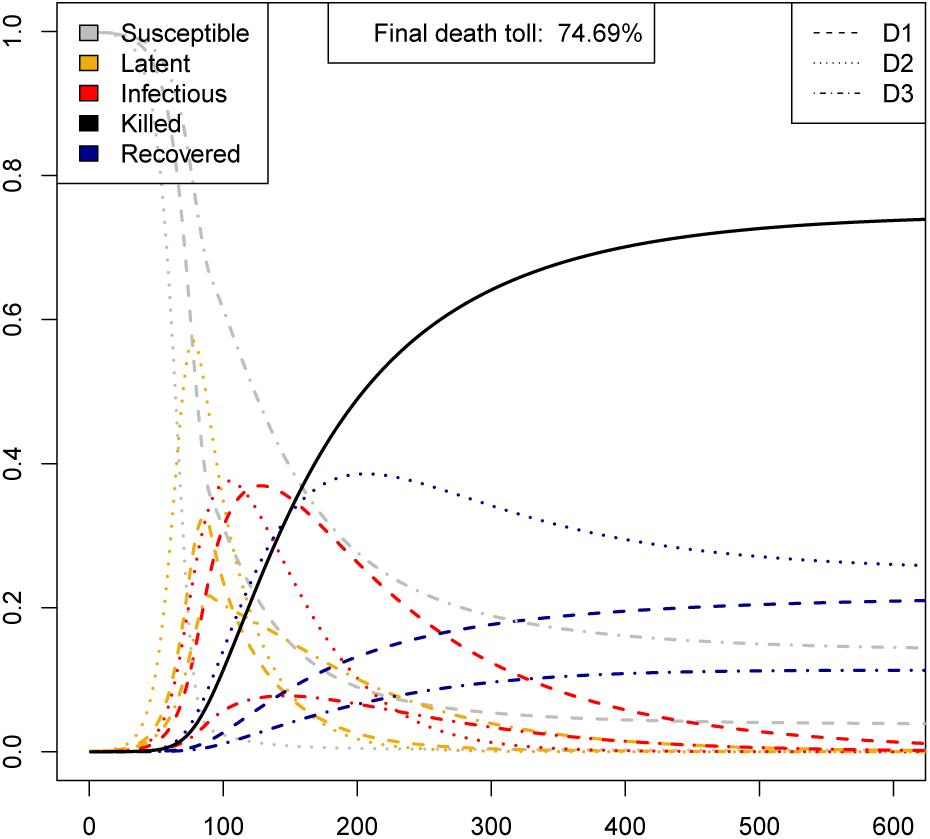
Highest Syndemic death rate com0 pared to worst individual disease case for 3 simultaneous outbreaks. The figure illustrates the spread of all three diseases when combined in a syndemic. Note that disease 3, which has the highest individual death rate, has the fewest infections during the syndemic because of its relatively slow spread. The other two diseases peak nearly simultaneously, well before the slower but higher fatality disease has a chance to spread more widely.

## 3, 4, and 5 Disease Pandemic Results

The simplified model used clearly shows that co-occurence of multiple diseases, without specific interactions or malevolent intent, are very unlikely to pose a much larger concern than a single disease from the set. The bottom line result about the increased danger of multidisease syndemics can be seen by noting two summary statistics from the analyses. First, the average death rate from 3, 4, and 5 disease syndemics was 44.9%, 47.9%, and 51.3%, respectively. This shows that multi-syndemics do become slightly worse when the number of diseases increases. However, the worst-case event among the 3, 4, or 5 diseases comprising the syndemics had averages of 46.5%, 49.3%, and 52.1%. This strongly implies that the primary driver of the increase is not a worse interaction among the diseases, but simply the fact that the greater number of competing diseases leads to an expected worse outcome. (Detailed results are plotted in Appendix B. All model and data files are available on the Github repository.)

It is still an open question how likely competitive versus interfering interactions among the diseases would be, and this could potentially significantly change the results. Other key uncertainties include difficult to mitigate diseases, and those where distancing and response measures are ineffective, though these issues are not specific to multipandemics.

## Further work

The model itself is somewhat rudimentary, and is much more useful as an indicative or conceptual tool for understanding how population reactions can change the dynamics of diseases. There are several ways in which the model could be extended, some more useful than others. One potential modification mentioned above include nonlinear reponse. If data were available to better understand the speed and form of distancing behavior, this could be an important extension.

Properly accounting for geographic heterogeneity would allow the model to consider border closures and screening, at significant computational expense. This may be particularly worthwhile for the single-disease case, to explore the potential impact of the delayed spread across borders and how it interacts with distancing and public health reaction.

Extending the multi-disease syndemic model to allow for disease interactions would allow much broader application, and while programmatically complex, does not necessarily add much computational complexity. More critically, such an extension would be useful to consider the actual relationships between specific potential diseases that might be implicated in an outbreak. In order to do so, however it may be necessary to find reasonable distributions for various cases in order to represent them. This would require more specific epidemiological research, such as the recent work by Adalja et al. [1], as well as difficult but important ongoing work on the estimation of the likelihood of future classes of disease [14].

Other extensions that would allow better representation requires adding compartments to the model. For instance, having multiple susceptible compartments with different distancing could allow representation of distancing fatigue. People would be hyper-vigilant for a time, then revert to being less strict. Having more than one infectious compartment would allow the representation of stealth diseases. The addition of compartments would be straightforward in the single disease case, but would make the multipandemic model even less computationally tractable, as should be clear from Appendix A.

## Conclusions

This analysis was intended to note an important likely feature of GCBR risk that was heretofore somewhat unclear; population reaction is a critical consideration when looking at worst case diseases, and is likely to drastically change the course of such diseases. The simplified model proposed is able to represent this, and show that most cases are self-limiting, and help identify cases where self-limiting spread will not occur; robust diseases, whihc spread despite public health reaction and distancing, acute diseases, which spread too quickly for such effects to make a significant difference, and stealth diseases, which are not noticed in time to prevent spread.

The model does not allow representation of specific parts of changing transmission dynamics. Despite this, for GCBR and otehr extreme risks, the models presented illustrate why population response and related factors that slow disease spread can be critical, sometimes even more than the characteristics of the disease alone. The model also shows the importance of better understanding how populations might react to these worst-case diseases, and implies that bio-surveillance able to give early warning, and effective post-outbreak warning and communications systems, are critical GBCR mitigation tools.

## Appendix A - Pictoral Guide to Multipandemic Complexity

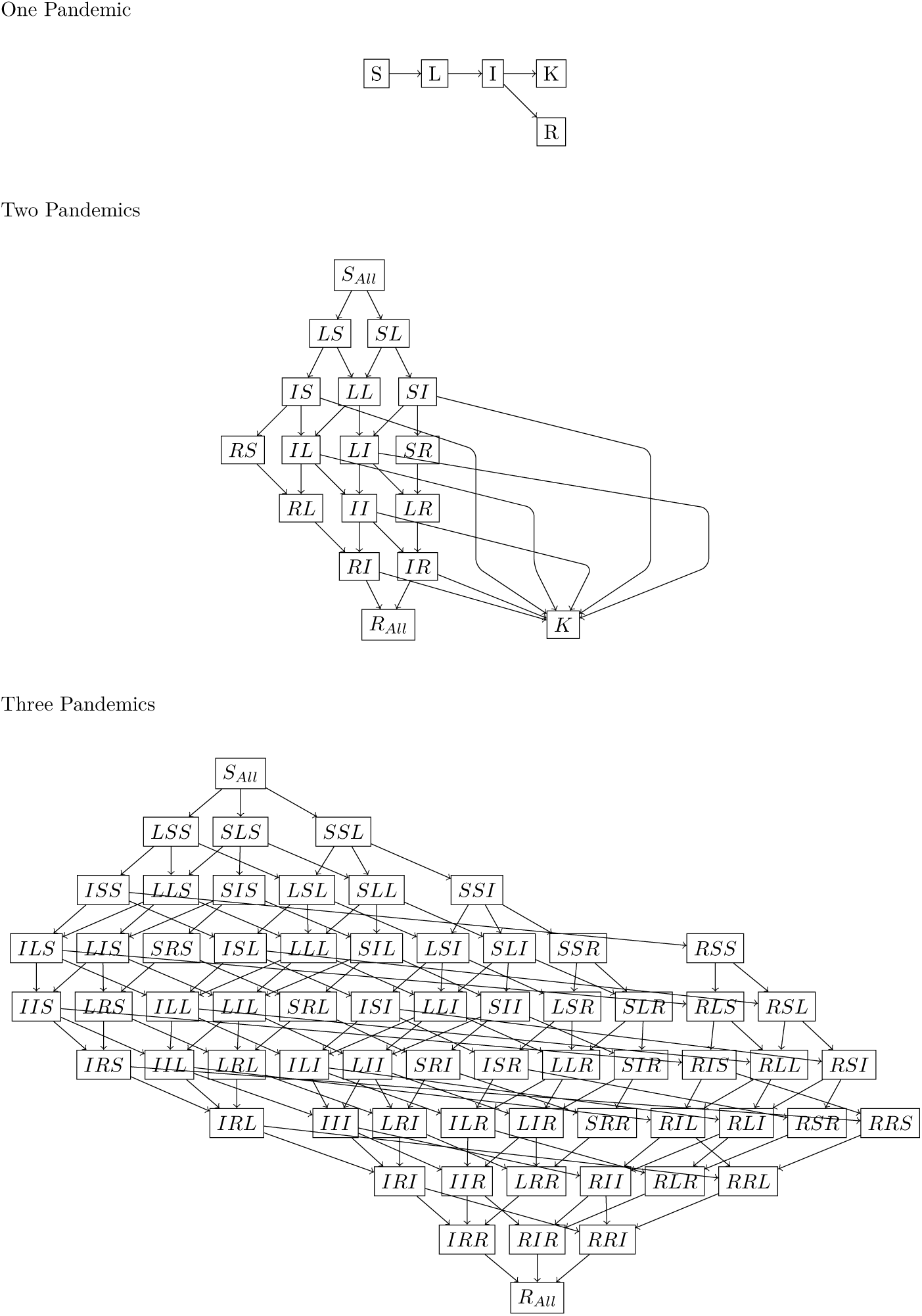

## Appendix B - Multipandemic Model Results

**Figure 5.**
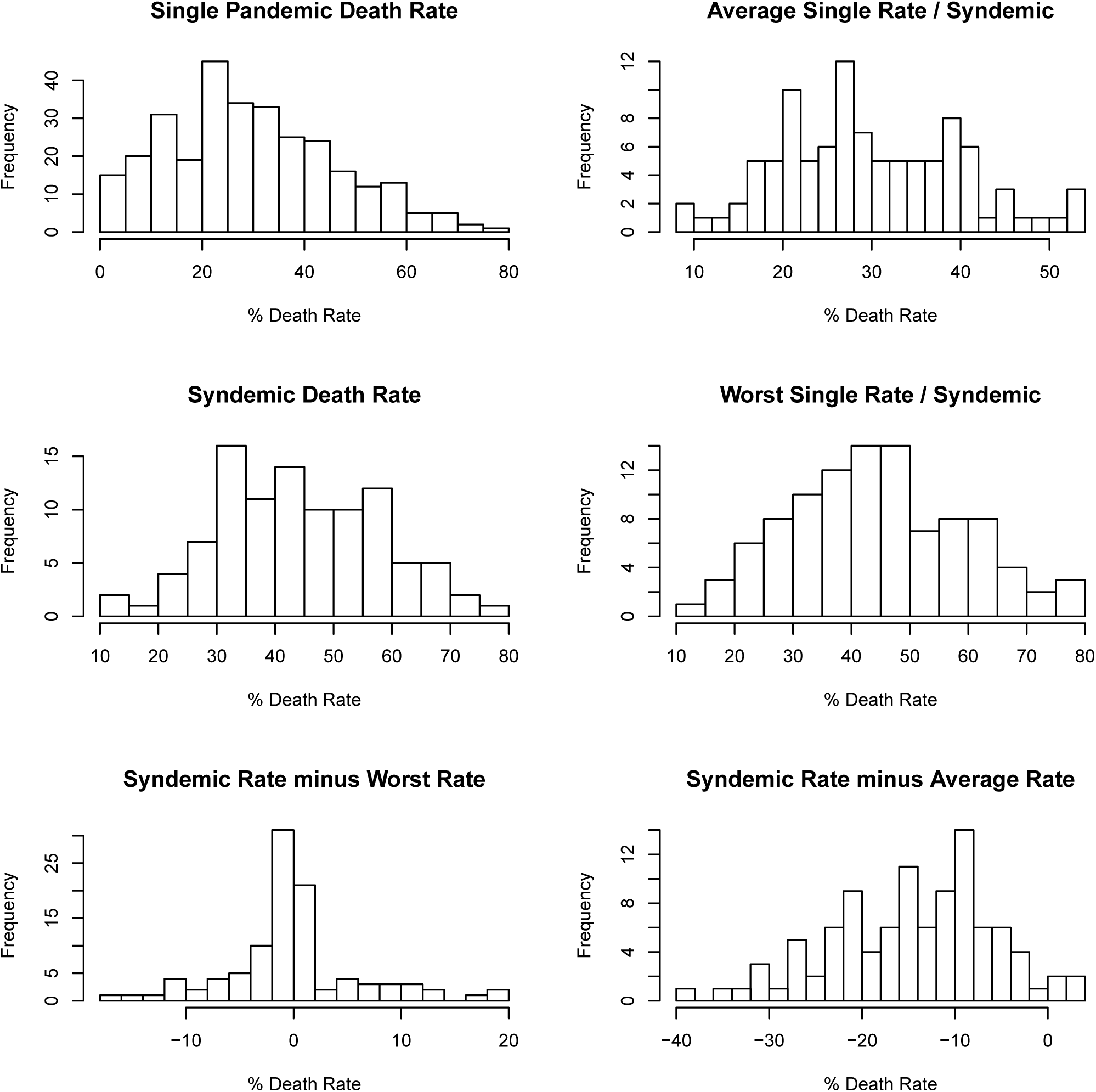
Syndemic summary histograms for 3 simultaneous outbreaks.

**Figure 6.**
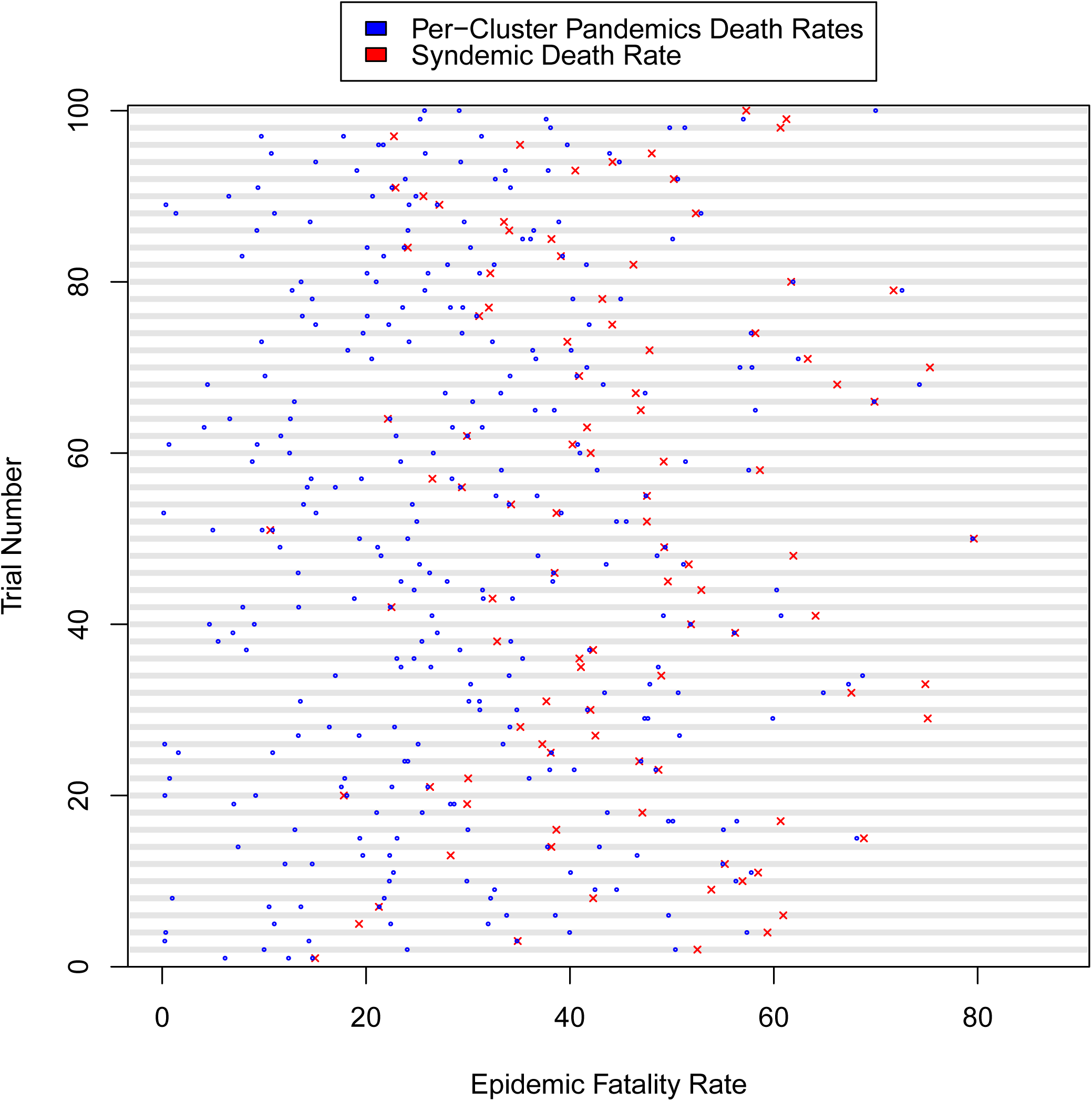
First 100 syndemic simulation epidemic population fatality rates, for 3 simultaneous outbreaks.

**Figure 7.**
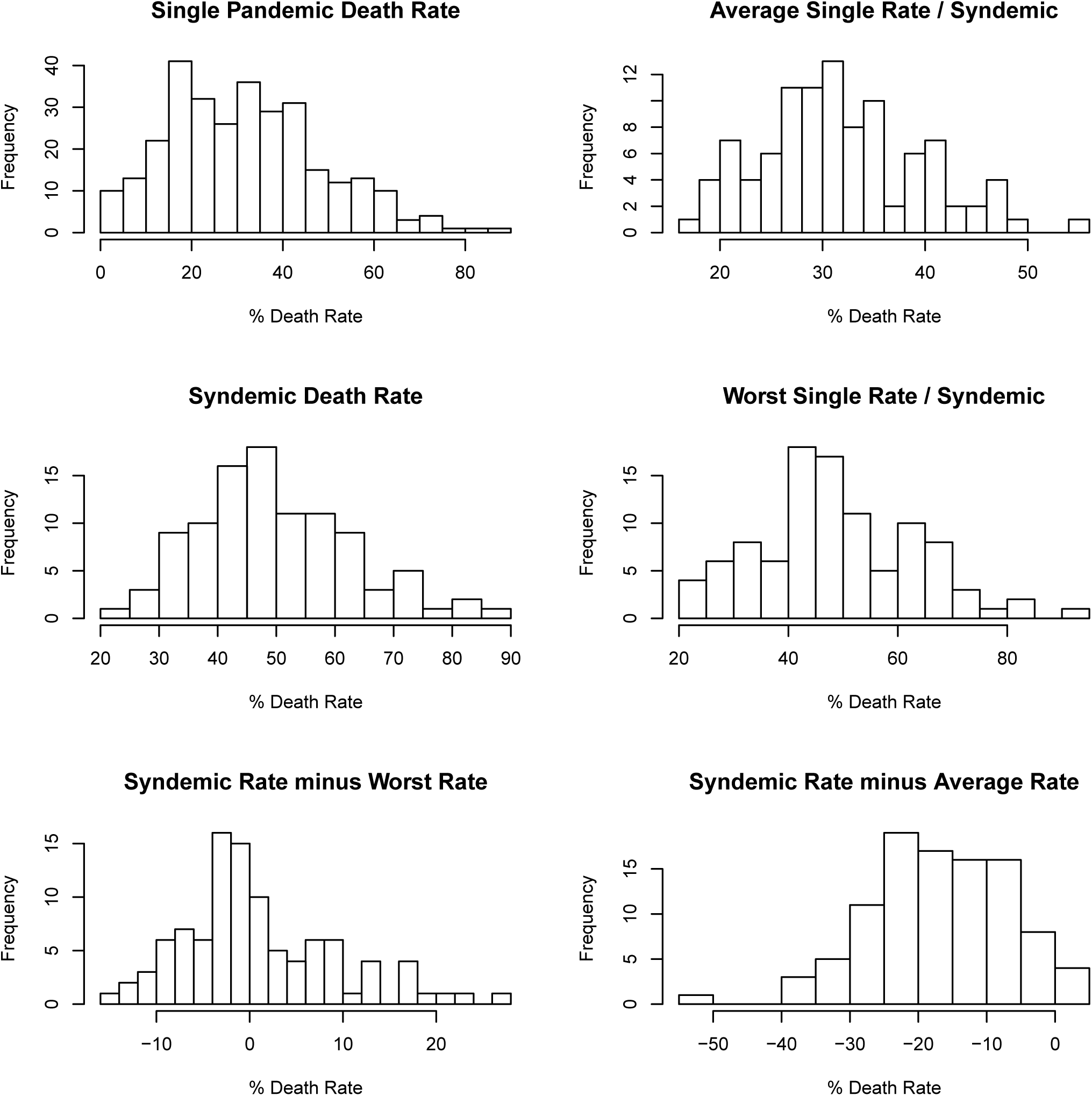
Syndemic summary histograms for 4 simultaneous outbreaks.

**Figure 8.**
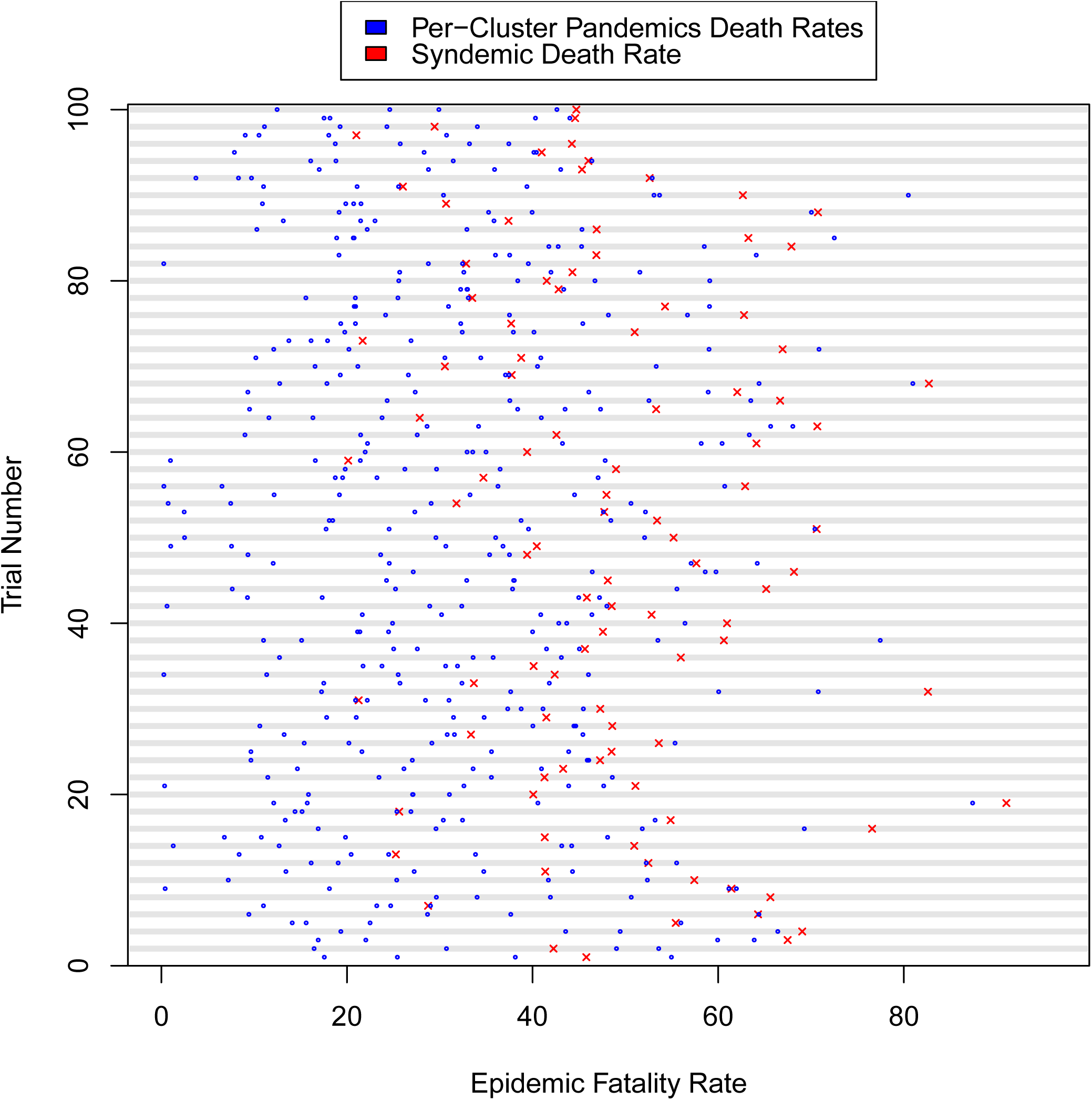
First 100 syndemic simulation epidemic population fatality rates, for 4 simultaneous outbreaks.

**Figure 9.**
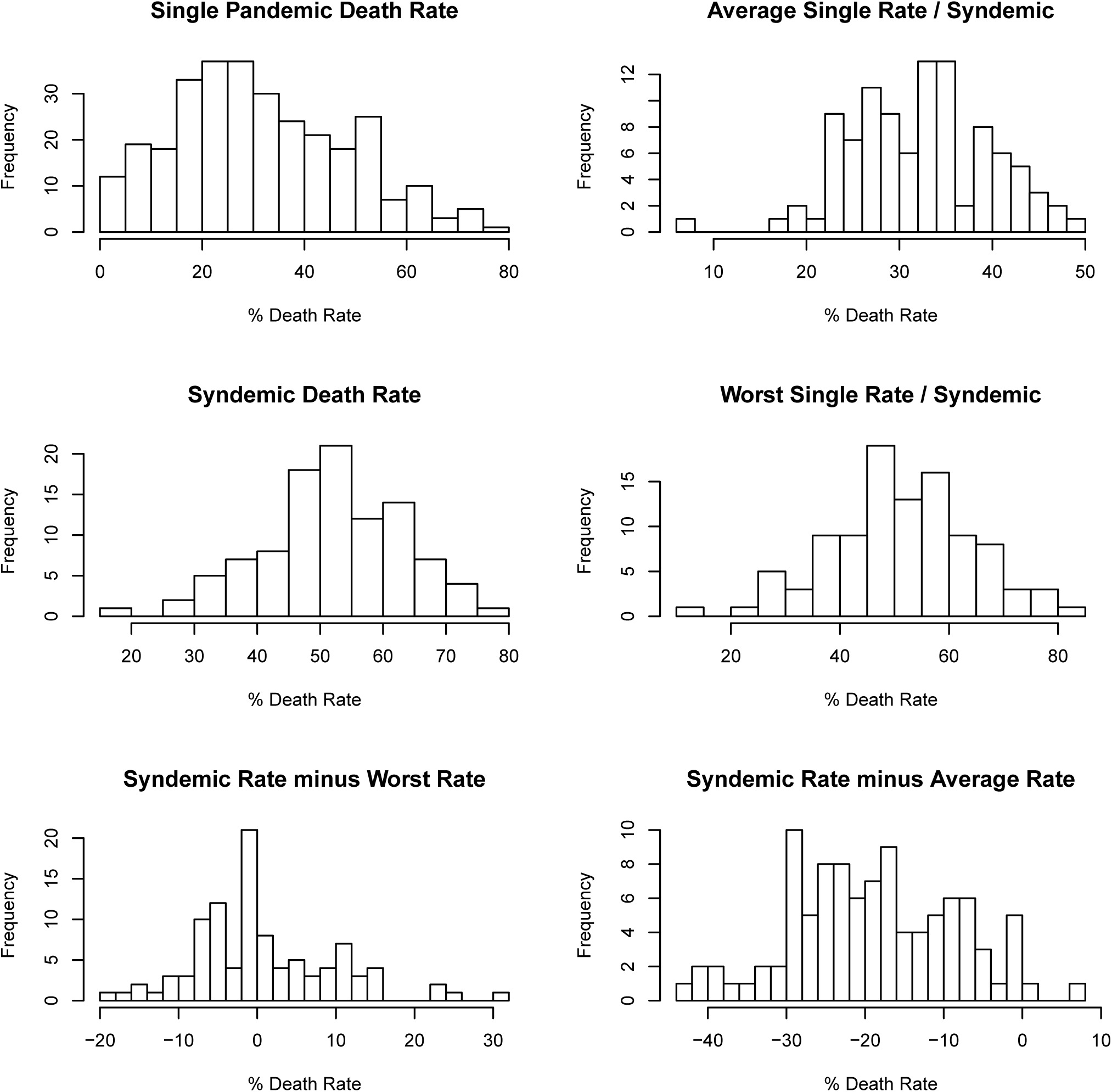
Syndemic summary histograms for 5 simultaneous outbreaks.

**Figure 10.**
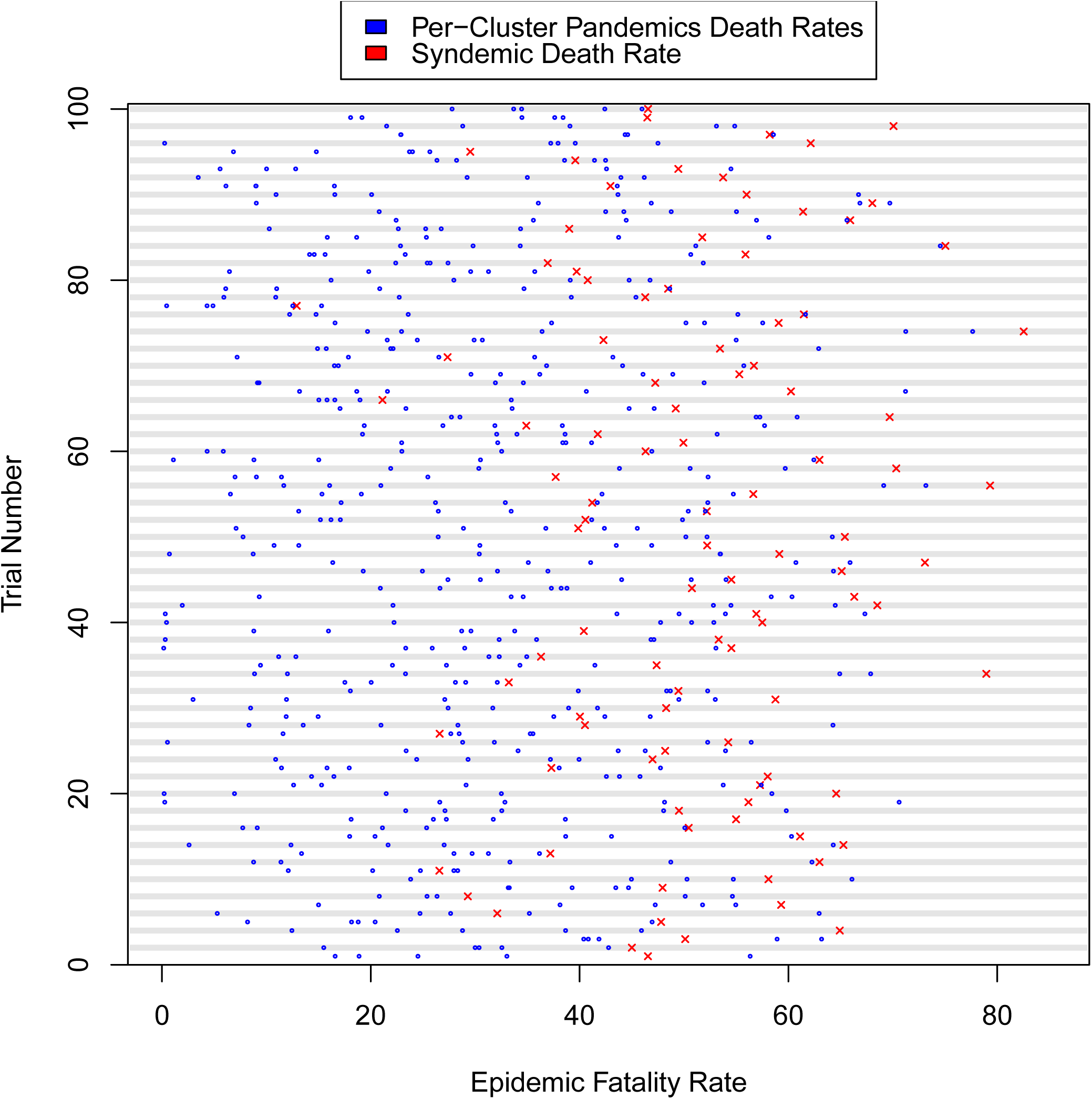
All 100 syndemic simulation epidemic population fatality rates, for 5 simultaneous outbreaks.

## Footnotes

1 Due to computational constraints, the 4- and 5-model versions were not run as many times. See Appendix A for a visual explanation.

## Acknowledgements

This paper is based on work done while visiting University of Oxford’s Future of Humanity Institute. I would like to thank them for the invitation, the incredible atmosphere, and the always-interesting discussion. Thanks to the Bio-risk team, especially Andrew Snyder-Beattie for insight and motivation, and Georgia Ray for her input and background research on syndemics. I would also like to thank Open Philanthropy, especially Claire Zabel, for funding the original research and the visit. Thanks also to Mike Bonsal, Gregory Lewis, and Raffaele Vardavas for their feedback.

## References

[1] AA Adalja, Watson M, Toner ES, Cicero A, and Inglesby TV. The Characteristics of Pandemic Pathogens. Technical report, Center for Health Security, 2018.

[2] Roy M Anderson and Robert M May. Infectious diseases of humans: dynamics and control. Oxford university press, 1992.

[3] Ben S Cooper, Richard J Pitman, W John Edmunds, and Nigel J Gay. Delaying the international spread of pandemic influenza. PLoS medicine, 3(6):e212, 2006.

[4] Neil Ferguson. Capturing human behaviour. Nature, 446(7137):733, 2007.

[5] Neil M Ferguson, Derek A T Cummings, Christophe Fraser, James C Cajka, Philip C Cooley, and Donald S Burke. Strategies for mitigating an influenza pandemic. Nature, 442(7101):448, 2006.

[6] Sebastian Funk, Erez Gilad, Chris Watkins, and Vincent A A Jansen. The spread of awareness and its impact on epidemic outbreaks. Proceedings of the National Academy of Sciences, 106(16):6872 LP – 6877, apr 2009.

[7] Robert J Glass, Laura M Glass, Walter E Beyeler, and H Jason Min. Targeted social distancing design for pandemic influenza. Emerging infectious diseases, 12(11):1671–1681, 2006.

[8] Andrea L Graham. Ecological rules governing helminthmicroparasite coinfection. Proceedings of the National Academy of Sciences, 105(2):566–570, 2008.

[9] R M Greer, P McErlean, K E Arden, C E Faux, A Nitsche, S B Lambert, M D Nissen, T P Sloots, and I M Mackay. Do rhinoviruses reduce the probability of viral co-detection during acute respiratory tract infections? Journal of Clinical Virology, 45(1):10–15, 2009.

[10] A Huppert and G Katriel. Mathematical modelling and prediction in infectious disease epidemiology. Clinical Microbiology and Infection, 19(11):999–1005, 2013.

[11] Istvan Z Kiss, Jackie Cassell, Mario Recker, and Pèter L Simon. The impact of information transmission on epidemic outbreaks. Mathematical biosciences, 225(1):1–10, 2010.

[12] Jeremy A Lauer, Klaus Röhrich, Harald Wirth, Claude Charette, Steve Gribble, and Christopher J L Murray. PopMod: a longitudinal population model with two interacting disease states. Cost effectiveness and resource allocation, 1(1):6, 2003.

[13] David Manheim. Eliciting Evaluations of Existential Risk from Infectious Disease, 2018.

[14] David Manheim. Questioning Overconfident Estimates of Risks from Natural Pandemics. Health Security, 2018.

[15] David Manheim, Margaret Chamberlin, Osonde A. Osoba, Raffaele Vardavas, and Melinda Moore. Improving Decision Support for Infectious Disease Prevention and Control: Aligning Models and Other Tools with Policymakers’ Needs. Technical report, RAND Corporation, 2016.

[16] Denis Mollison. Epidemic models: their structure and relation to data, volume 5. Cambridge University Press, 1995.

[17] Brian Mustanski, Robert Garofalo, Amy Herrick, and Geri Donenberg. Psychosocial health problems increase risk for HIV among urban young men who have sex with men: preliminary evidence of a syndemic in need of attention. Annals of Behavioral Medicine, 34(1):37–45, 2007.

[18] Romualdo Pastor-Satorras, Claudio Castellano, Piet Van Mieghem, and Alessandro Vespignani. Epidemic processes in complex networks. Reviews of Modern Physics, 87(3):925–979, aug 2015.

[19] RStudio, Inc. Easy web applications in R., 2013. Annotation: URL: \url{http://www.rstudio.com/shiny/}

[20] Monica Schoch-Spana, Anita Cicero, Amesh Adalja, Gigi Gronvall, Tara Kirk Sell, Diane Meyer, Jennifer B Nuzzo, Sanjana Ravi, Matthew P Shearer, and Eric Toner. Global catastrophic biological risks: toward a working definition. Health security, 15(4):323–328, 2017.

[21] Linda A Selvey, Catarina Antao, and Robert Hall. Evaluation of border entry screening for infectious diseases in humans. Emerging infectious diseases, 21(2):197, 2015.

[22] Zhen Wang, Michael A Andrews, Zhi-Xi Wu, Lin Wang, and Chris T Bauch. Coupled diseasebehavior dynamics on complex networks: A review. Physics of Life Reviews, 15:1–29, 2015.

[23] Jaime Yassif. Reducing global catastrophic biological risks. Health security, 15(4):329–330, 2017.

